# Quantitative comparison of camera technologies for cost-effective Super-resolution Optical Fluctuation Imaging (SOFI)

**DOI:** 10.1101/413179

**Authors:** Robin Van den Eynde, Alice Sandmeyer, Wim Vandenberg, Sam Duwé, Wolfgang Hübner, Thomas Huser, Peter Dedecker, Marcel Müller

## Abstract

Super-Resolution (SR) fluorescence microscopy is typically carried out on high-end research microscopes. Super-resolution Optical Fluctuation Imaging (SOFI) is a fast SR technique capable of live-cell imaging, that is compatible with many wide-field microscope systems. However, especially when employing fluorescent proteins, a key part of the imaging system is a very sensitive and well calibrated camera sensor. The substantial costs of such systems preclude many research groups from employing super-resolution imaging techniques.

Here, we examine to what extent SOFI can be performed using a range of imaging hardware comprising different technologies and costs. In particular, we quantitatively compare the performance of an industry-grade CMOS camera to both state-of-the-art emCCD and sCMOS detectors, with SOFI-specific metrics. We show that SOFI data can be obtained using a cost-efficient industry-grade sensor, both on commercial and home-built microscope systems, though our analysis also readily exposes the merits of the per-pixel corrections performed in scientific cameras.

## 1 Introduction

The ability to quantitatively and specifically image biological structures of interest makes fluorescence microscopy an essential tool in the life sciences. However, its spatial resolution is limited in a conventional imaging system due to the diffraction of light. To overcome this limitation, a range of Super-Resolution (SR) techniques has been developed and enhanced over the last two decades^1,2^. These techniques offer a tremendous improvement in spatial resolution, though in most cases at a cost in temporal resolution. They also typically require high-end imaging systems, limiting their use to those scientists who have access to such equipment. Driven by this observation, several initiatives have been started over the past years to ‘democratize’ fluorescence microscopy. These aim for a careful trade-off between performance and cost, instead of maximizing system performance. This goal can be achieved by a combination of open-source / open-access software and hardware blueprints, as well as repurposing commodity industrial or consumer hardware^3–9^.

Highly sensitive cameras often pose a substantial portion of a microscope’s cost. A high-end sCMOS camera, for example, can exceed 10.000 Euro at the time of this writing. In comparison to conventional industry-grade cameras, priced under 1000 Euro, the value of these scientific-grade cameras lies in their higher performance in terms of Signal-to-Noise-Ratio (SNR), increased acquisition speed, higher detection sensitivity and extensive manufacturer-provided calibration. Nevertheless, recent work has shown that even within the demanding realm of SR microscopy, acceptable to good performance can be obtained with industry-grade components. For example, an industry-grade camera is sufficient for super-resolution techniques based on single molecule localization microscopy (SMLM) such as dSTORM and fPALM, which was confirmed by extensive analysis and characterization of these camera systems^5,10^. A key reason for this success is the fact that most organic fluorophores are very bright, emitting many photons before photo-destruction occurs, which reduces the overall sensitivity requirements for the camera. Furthermore, the magnification in these systems is typically tuned so that the point-spread function is sampled by multiple detector pixels at once, which can reduce the impact of imperfect pixelwise calibration. While localization microscopy provides a very high spatial resolution it does so at a cost to temporal resolution, with the labeling strategies leading to the highest resolutions even requiring fixation of the sample. However, a vast amount of biological questions requires the use of SR live-cell imaging techniques. It is therefore important to question to what extent the inexpensive detectors can also be employed for this purpose.

Super-resolution Optical Fluctuation Imaging (SOFI)^11^ is an established super-resolution microscopy method, working not only with quantum dots^11^ and organic dyes^12^, but also with genetically-encoded fluorescent proteins^13,14^, which is a great asset for biological research. This technique combines an isotropic increase in spatial resolution with a good temporal resolution, rendering it especially useful for living biological samples^13–17^. Like other diffraction-unlimited techniques, such as SMLM, the key ingredient for SOFI microscopy is the use of dynamic fluorescent labels. SOFI relies on the statistical analysis of multiple images acquired from the same sample, labelled with fluorophores that show independent and transient non-emissive intermittencies, or ‘blinking’ behavior. Probes that display these dynamics, such as photochromic fluorescent proteins, have been shown to result in a two-to three-fold spatial resolution improvement over the classical diffraction-limited resolution^13,14^. The SOFI framework even allows for multiplexing of several emitters, with highly similar steady-state fluorescence characteristics, using differences in their blinking behavior^18^. In addition to the conventional visualization of fluorophore distributions, SOFI has also been used to visualize biosensor activities with similar spatial resolution enhancements^19,20^.

In contrast to SMLM, however, the key of SOFI-based analysis is its reliance on spatio-temporal correlations in the signal. To avoid artifacts, this places rather unusual demands on the image sensor, as the acquisition system itself should not introduce correlations (stemming i.e. from read-out electronics) into the raw images. In this work we set out to establish and characterize to what extent different camera architectures can be used for SOFI, investigating both an inexpensive industry-grade camera and high-end scientific cameras. We focused on the use of genetically-encoded fluorescent proteins, whose overall brightness is considerably lower than those of organic dyes. We found that SOFI imaging can be performed well with all three systems, providing good baseline imaging with the industry-grade camera, though the scientific systems provide increased quantitative accuracy and higher sensitivity in the red region of the visual spectrum. Our work directly enhances the accessibility to the scientific community, and readily enhances the utility of this versatile imaging technique. We additionally demonstrate its use on a home-built, free-standing microscope, in line with the goal of reducing the overall system costs.

## 2 Results

We directly compared three different cameras via pairwise evaluation through a 50/50 beam splitting cube (see Table 1 and Figure 1, panels C & E). This approach allows the same signal (frame by frame image of the sample) to be detected by two cameras, and thus the quantitative analysis is not influenced by sample variation. For comparison we selected two high-end cameras, commonly employed for super-resolution imaging, with two different sensor architectures: the Hamamatsu Fusion (sCMOS) and the Andor Ixon-ultra-897 (EMCCD). As the third camera we employed an IDS μEye UI-3060CP-M-GL Rev.2 industry-grade CMOS camera (list price around 650 Euro). This latter camera has been represented, with a similar purpose, as a viable alternative for dSTORM imaging^5^.

**Figure 1.:**
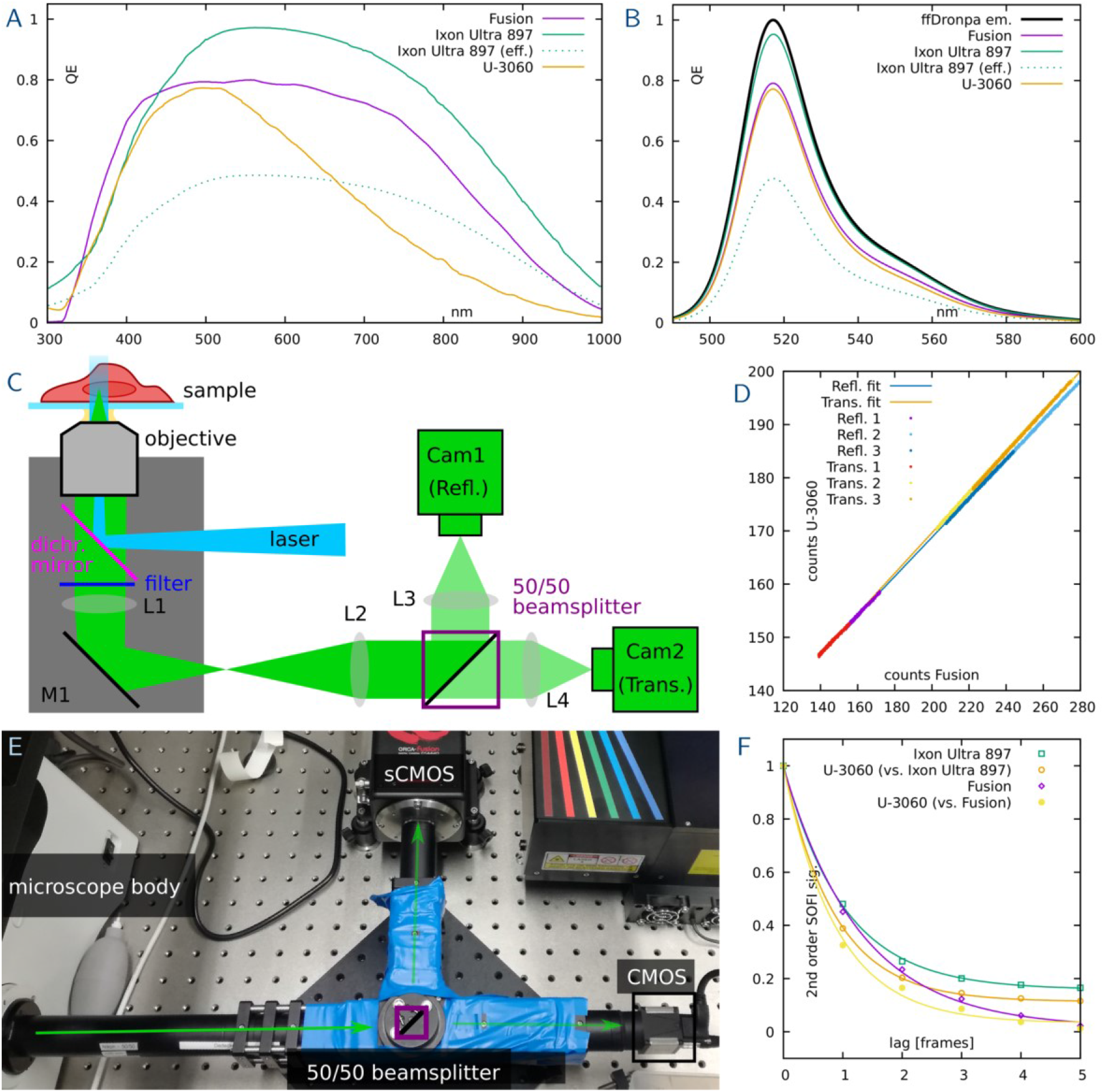
Quantum efficiency and calibration of the SOFI experiments. The raw quantum efficiency curves provided by the manufacturers (panel A) show a high sensitivity for green light. To obtain the detection efficiency for ffDronpa, these curves have been weighted with its emission spectrum^31,32^ (panel B) and it becomes obvious all cameras should be well suited to detect ffDronpa. For the emCCD, an effective QE, weighted with a factor of 2, is also displayed to account for its readout process^23^. A sketch (panel C) and photograph (panel E) of the beamsplitter arrangement illustrated the setup in use. L1 is part of the microscope body (grey box) and is normally used to focus on a camera. Here, L2 (f=125mm) is utilized to parallelize the detection beam which is needed to avoid aberrations with the 50/50 beam splitter. The blue tape is used to make the construction fully light-tight. The splitting ratio has been tested by swapping cameras between the reflective and transmissive configuration for three samples each and comparing their count slopes (panel D). Slope fitting yields an effective splitting ratio of 49% to 51% in the beam splitting device, which is well within expectation and tolerance for the experiments. Lastly, excitation light levels were tuned so that a τ of around 1 was reached for the 40ms exposure times used in the experiments (panel F).

**Table 1.:**
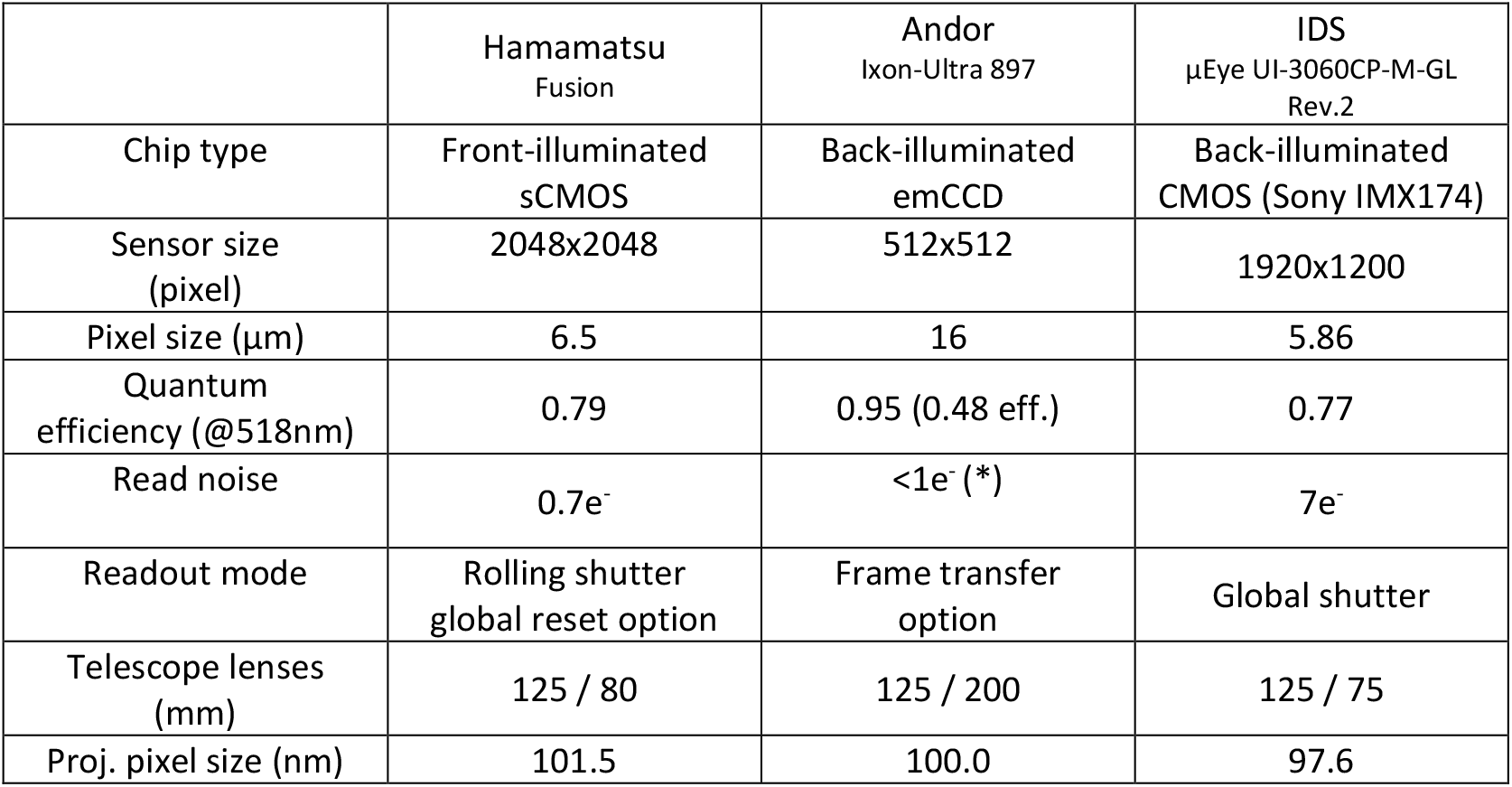
Technical comparison of the three cameras tested, and focal length used in the 50/50 beam splitter. The latter were selected such that the overall magnification for each camera (using a 100x objective, and re-magnifying in the relay telescope) yields closely matching projected pixel sizes in the range of 100nm. (*) The read noise of an emCCD camera is a function of the em-gain; for the Ixon Ultra, it is specified to 50e^−^ (rms), thus reduced to < 1e^−^ by an em-gain of 50x or more (see manufacturer data sheet).

### 2.1 Sample and tuning of data acquisition

To enable a critical assessment of the camera SNRs and their sensitivity, we used Cos-7 cells where microtubules were fluorescently stained (see Methods). These fiber-like, three-dimensional cytoskeletal structures show a wide variety of thicknesses and exhibit branching, which makes them excellent structures to demonstrate super-resolution imaging. We stained these structures with a photo-switching protein, photochromic ffDronpa^21^, which was linked to microtubule-associated protein 4 (MAP4). The emission spectrum of ffDronpa peaks at about 515 nm, which coincides well with the region of highest detection efficiency of all three cameras (see Figure 1, panels A & B). In all cases we adjusted the excitation intensity to obtain an emitter τ-value^18^ (decorrelation time) of around one exposure time (see Figure 1, panel F). This value offers the highest signal for reconstruction using conventional second-order SOFI imaging without lag time.

### 2.2 Scientific microscope system

For the camera comparison we started off with a high-end research microscope. We used a *Nikon Eclipse TI2* microscope body outfitted with a high-power 200 mW 488 nm laser, coupled to the microscope through a fiber illuminator and a 100x oil-immersion objective lens (see method section for component details). This setup is in routine use for high-end super-resolution imaging. A 50/50 beam splitter (see method section and Figure 1, panels C & E) was inserted, followed by the detectors and their dedicated tube lenses. These lenses varied in focal length to obtain a similar projected pixel size (97 – 102 nm with a 100x objective lens and Nikon tube lens) for a fair comparison between cameras (see Table 1). The split ratio between the transmission and reflection arm of the beam splitter was verified (see method section and Figure 1, panel D) and found to not introduce a mismatch of more than 1%. The cameras were set to an exposure time of 40 ms and were electronically synchronized (see methods) to both minimize sample bleaching and ensure frame-by-frame comparable data acquisition.

We processed the acquired data sets with the SOFI algorithm. For this the open-source Localizer package^22^, with a user interface and analysis framework in the Igor Pro software package (Wavemetrics, Portland, USA) was used. An image stack of 250 frames proved sufficient to clearly showcase the resolution improvement achieved by SOFI and to compare the camera systems both in terms of visual image quality (see Figures 2 and 3, panels A to C each) and quantitatively (see Figures 2 and 3, panels D to F each). We chose Fourier Ring Correlation (FRC) analysis, SNR and a SOFI-specific spatial correlation analysis for extracting the PSF shape (PSF) as suitable metrics for the comparison (descriptions of applying FRC to SOFI data and the spatial correlation analysis are provided in methods).

**Figure 2.:**
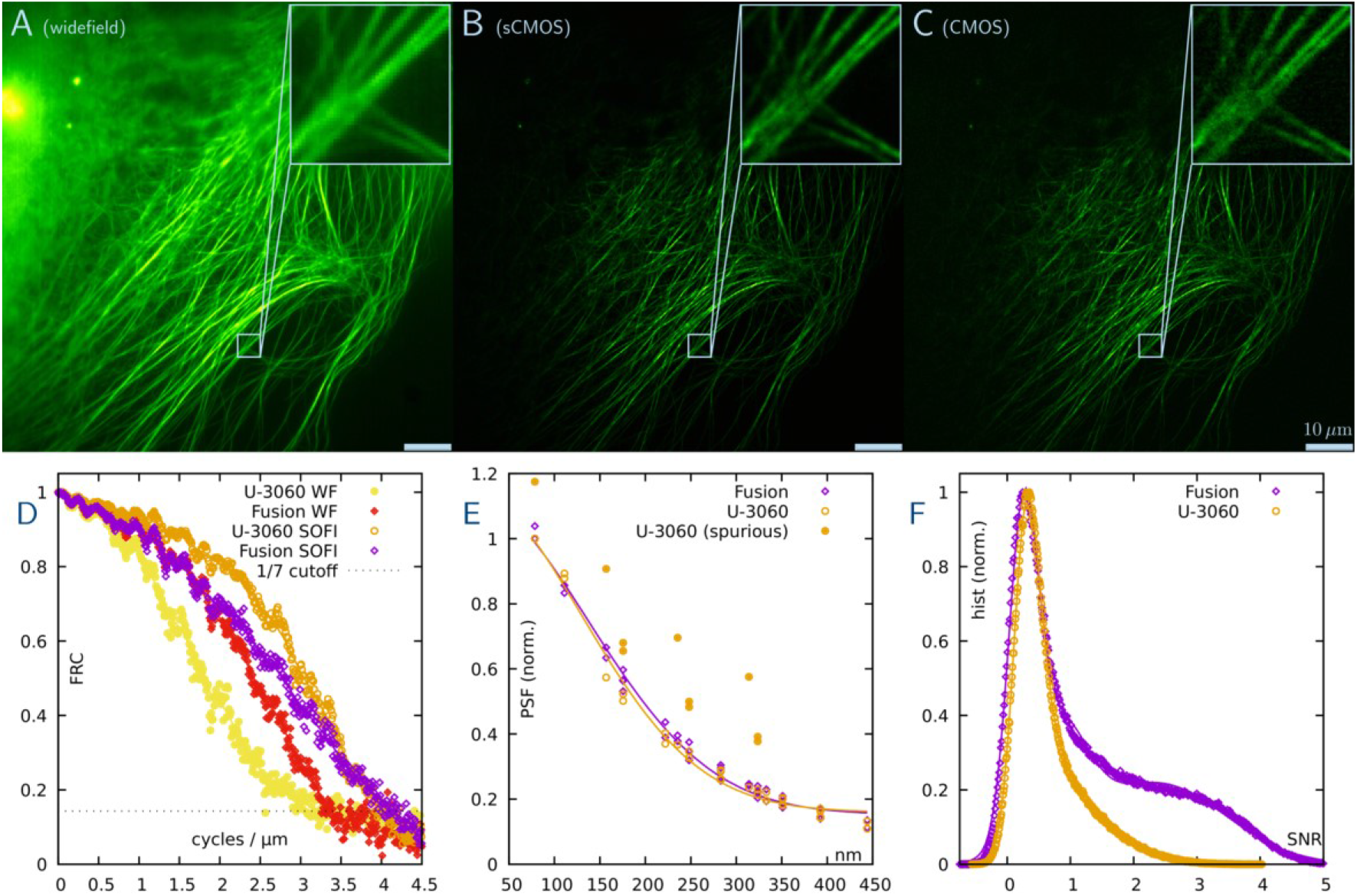
SOFI imaging comparing the sCMOS and the CMOS camera. The microtubules of the Cos-7 cells were stained with ffDronpa. The widefield image (panel A) is generated as the average of 250 analyzed SOFI frames (stack subsamples in 5x 50 blocks each, see FRC methods section). The upper left region shows lots of out-of-focus light as blur, the lower right part features structures of interest. The SOFI reconstruction of the sCMOS camera (panel B) both removes the out-of-focus contribution, and enhances resolution so fibers can be separated much more clearly. The SOFI reconstruction of the CMOS camera yields comparable results (panel C), whilst also showing higher noise levels, which is in line with expectations. The resolution, measured via Fourier ring correlation (see methods), reflect this (panel D), where the CMOS camera picks up more noise both in the wide-field and in the SOFI analysis. Similarly, the signal-to-noise ratio is higher for the sCMOS camera (panel F). Importantly, cross-pixel correlations, which are a typical artifact of CMOS technology, are unsurprisingly present in the CMOS datasets, but seem heavily reduced in the current-generation sCMOS chips tested here (panel E). Scalebar 10 μm, insets 5 μm.

**Figure 3.:**
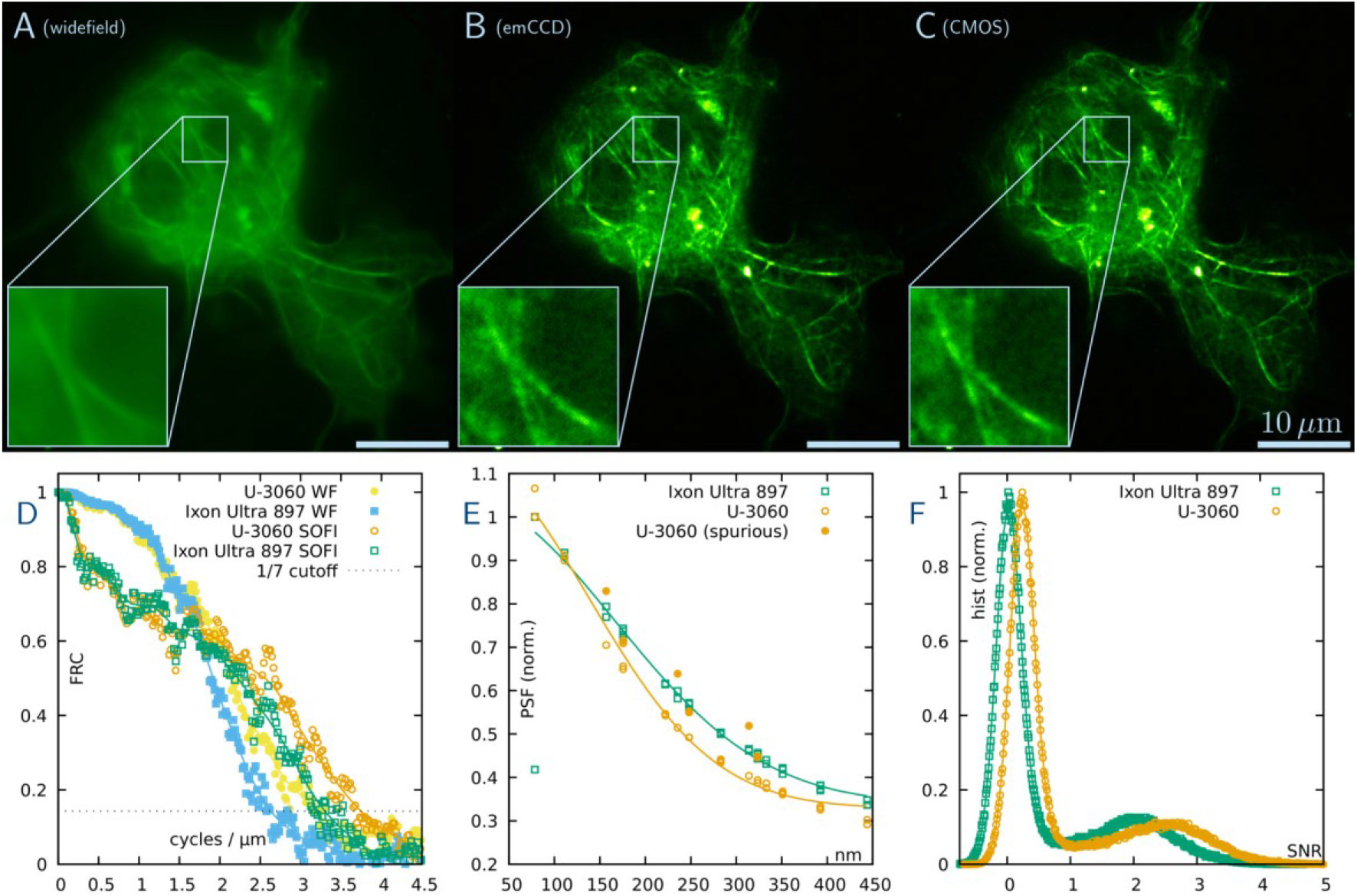
SOFI imaging comparing the emCCD and the CMOS camera. Again, microtubules in Cos-7 cells were labelled with ffDronpa. The widefield image (panel A) is generated as the average of 250 analyzed SOFI frames (stack subsamples in 5x 50 blocks each, see FRC methods section). The SOFI reconstruction of the emCCD camera (panel B) enhances resolution so two fibers not distinguishable in wide-field can be separated (see magnified insets). The SOFI image obtained via the CMOS camera yields the same improvement, with arguable higher contrast (panel C). To understand this performance difference, it is important to know that the pure photon counts reached in SOFI experiments heavily favors CMOS technology (see main text discussion). This also reflects in the CMOS outperforming the emCCD slightly in both FRC resolution (panel D) and signal-to-noise ratio (panel F). However, the emCCD does not show spurious pixel correlations typical for CMOS technology (panel E). Scale bars 10 μm, insets 5 μm.

We find that the current-generation sCMOS camera slightly outperforms the CMOS camera in the achieved FRC resolution and provides a much higher SNR, with the SNR peaks in Figure 2 being 1.88x higher for the sCMOS compared to the CMOS camera. Also, no additional correlations stemming from the camera are detected in the sCMOS camera, while clear spurious correlations are observed in the CMOS which will serve as a (limited) source of bias in the SOFI imaging (see figure 2 panel E). Visually, the SOFI reconstructions obtained with both cameras provide a clear resolution improvement and background reduction over the wide-field images, while the CMOS data arguably appears somewhat noisier and grainier.

Comparing the emCCD and the CMOS system, we find that, surprisingly, both cameras perform much more similar, and that the CMOS camera slightly outperforms the emCCD system, both in the achieved SNR as in the FRC resolution reached. However, the emCCD system does not show spurious correlations in the extracted PSFs, which the CMOS system continues to introduce. The results might seem surprising, given that emCCD systems are typically viewed as very sensitive, but are explained by the average amount of photons observed per pixel, and the resulting read noise and shot noise statistics (see discussion).

### 2.3 Home-built microscope system

Home-built systems offer a combination of flexibility and lower cost compared to commercial systems, and therefore present attractive avenues for reducing the overall cost of super-resolution imaging. We tested the viability of such a system, build from free-standing optics, for SOFI measurements with the industry-grade CMOS camera. The system is comprised of a 50 mW 473 nm diode laser and a 60x oil immersion objective in epi-fluorescence configuration. A 250 mm tube lens was used to obtain a projected pixel size of 70 nm (see figure 4, panel A and B). At a given resolution limit of about 195 nm, this slightly oversamples the detection. However, since the field of view of the CMOS chip is very large, this configuration was preferred to allow for a telecentric 2f configuration. While more limited in operation (no motorized stage, no automated opto-mechanics) and flexibility (only two fixed laser excitation lines in the case of this system), the image quality is comparable to a research microscope (see figure 4, panel C, D and E). Overall, this system can be bought and self-assembled at a total cost of about 16.500 Euro, which is much lower than the cost of typical SR instruments.

**Figure 4.:**
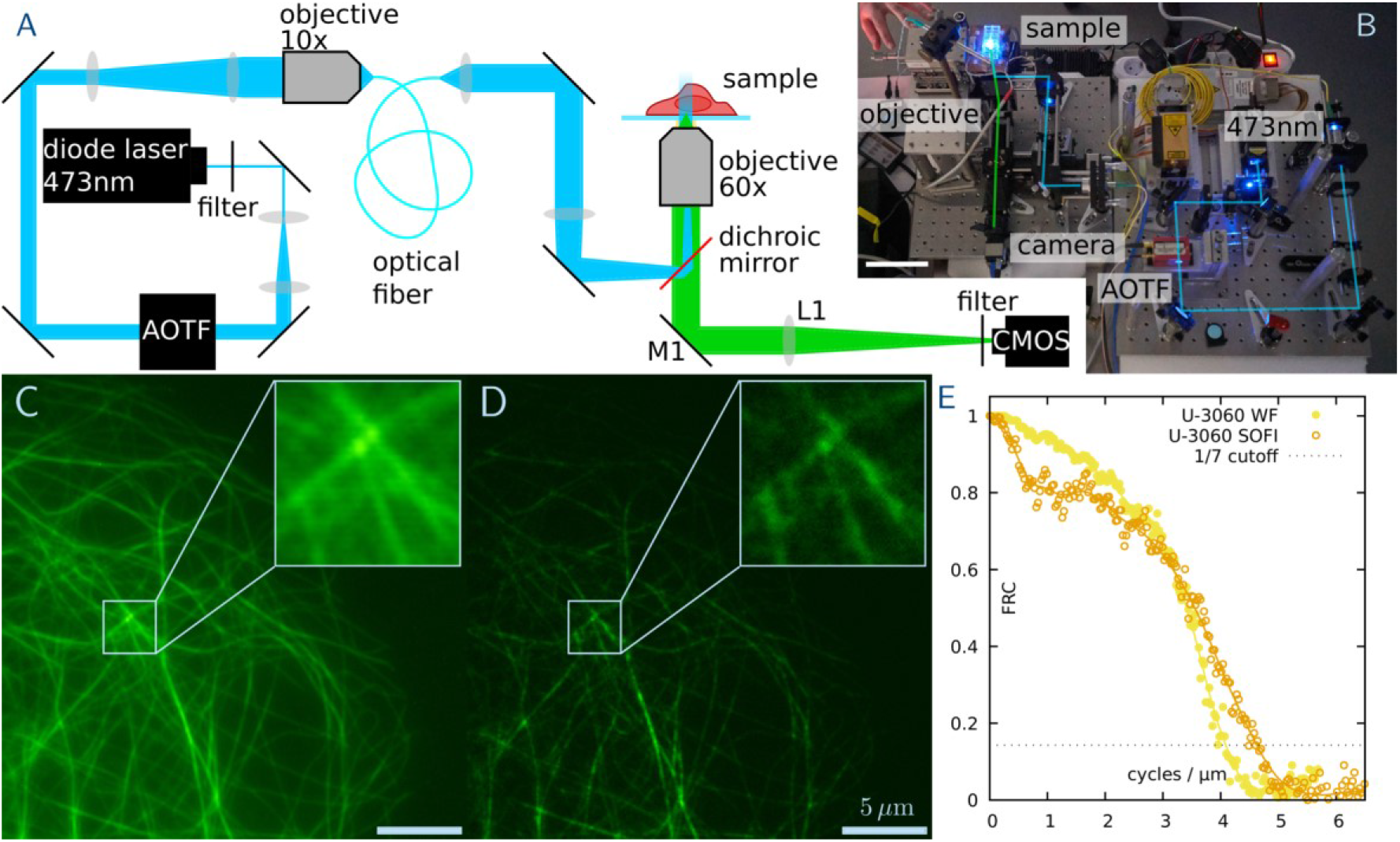
SOFI imaging on the home-built microscope system with the CMOS camera. (panel A) Schematic sketch of the system and (panel B) a photograph of the system during a measurement (for details see methods). Scale bar, 10 cm. (panel C) Average widefield image of microtubules in Cos-7 labelled with ffDronpa. (panel D) SOFI image of 250 reconstructed frames. Exposure time 10 ms. Scale bar 5 μm, inset 2 μm. A direct comparison shows clearly the resolution improvement (see insets) which is quantified by the FRC analysis (panel E).

The SOFI measurements performed on this system do not pass a beam-splitting arrangement, so all photons are collected on the same camera. This of course greatly improved the raw signal’s SNR, and thus the achieved FRC resolution of the system (as it relies on an SNR cutoff). However, using a single camera is of course closer to a ‘real world’ application of the system, and showcases the quality one could reach in SOFI imaging when employing this type of cost-efficient hardware.

The visual results (fig. 4 panel C and D) again show a clear reduction in background and an improvement in resolution when comparing wide-field and SOFI reconstruction. This is mirrored by the FRC data obtained for those images (fig. 4 panel E). Since we did not perform a camera comparison on this system, a SNR analysis is not meaningful, as it is highly sample-dependent (changing with staining efficiency, expression levels and such) and thus requires a direct, frame-by-frame comparison.

## 3 Discussion and conclusion

Our findings provide a quantitative comparison of both a state-of-the-art sCMOS and an established, high-end emCCD camera with a CMOS solution targeted at industrial applications, with metrics specific to SOFI super-resolution imaging.

The industry-grade CMOS camera is clearly capable of acquiring SOFI datasets of sufficient quality on realistic day-to-day samples, even when using fluorescent proteins as labels. In a direct comparison, its image quality is not on par with a current-generation sCMOS system. The images are visually noisier, and the associated SNR and FRC metrics (which implicitly includes an SNR-style weighting) reflect this. This is to be expected, as both cameras share the same sensor technology, but the ‘scientific’ CMOS implementation is much more refined in terms of read-out noise and calibration.

In direct comparison of the emCCD and the CMOS systems, both the visual image quality and the SNR and FRC metrics are almost on par. The CMOS camera even performs slightly better than the emCCD system, which might seem surprising, as the emCCD system is of course equally fine-tuned to scientific applications. However, the difference in sensor technology, especially the reliance on the electron-multiplication stage, puts the emCCD system at a disadvantage for the photon count levels used in SOFI imaging.

Looking at both these findings and the signal formation aspect of SOFI imaging, two technical characteristics seem to influence SOFI imaging quality the most: mainly the quantum efficiency of the sensor, and secondarily its read noise. The Sony sensor in use provides a high quantum efficiency in the green, with a pronounced fall-off towards red and infrared light (see figure 1 panel A). As the sensor is marketed for a broad range of applications, this is a sensible optimization by the manufacturer. For the application presented here, this aligns well with the green (peak at 515 nm) emission spectrum of ffDronpa (see figure 1 panel B) and yields a detection efficiency almost identical to the sCMOS camera system (for red and far-red emitting dyes, as often used in dSTORM microscopy, there would be a significant difference in detection efficiency). The difference between the sensors manifests in their read noise characteristics, where the sCMOS camera yields 0.7e- to 1.4e- (depending on readout speed) compared to 7e- for the Sony sensor. We expect that a typical ‘blink’ of ffDronpa will a few hundred photons distributed over the pixels sampling its PSF, so this change in read noise is likely the main reason for the change in SOFI SNR.

The emCCD sensor provides an even higher quantum efficiency over the full spectral range, but at these typical SOFI photon count levels, the excess read-noise introduced by the electron multiplying stage cannot be compensated by the higher quantum efficiency (our plots in figure 1 panels A & B include the typical^23^ 2*x* weighted quantum efficiency plot for the emCCD chip to account for this effect). This effect is documented in comparisons of sCMOS and emCCD technologies, where emCCD systems for some time are only recommended for the lowest photon count levels, and seems to hold even for current-generation, industry-grade CMOS technology.

Lastly, sensor readout uniformity, linearity and ‘spurious correlation’ influence the quality of a SOFI image. While uniformity and linearity are already well documented^5^, the effect of inter-pixel correlations becomes very apparent in SOFI imaging: If a signal, typically a high photon count, spatially or temporally influences the count rate in neighboring pixels, this effect shows up as an unreasonably high correlation in the PSF-estimate plots (panel E in figure 2 and 3). Causes for these correlations can be both non-uniform pixel cross-talk and latent effects in the read-out electronics. Here, both the emCCD and the sCMOS cameras seem to be highly calibrated and optimized, both showing no spurious correlations, while the industry-grade CMOS system shows a moderate amount of correlation not stemming from single molecule blinking. While these signals become apparent in an in-depth quantitative analysis, they still seem to be controlled enough to not have an adverse effect on image quality.

In conclusion, we could demonstrate that a current generation industry-grade CMOS camera system can acquire SOFI raw data with enough sensitivity and fidelity to yield convincing image quality on typical biological samples. A current-generation sCMOS system provides enhanced sensitivity in the red range of the spectrum, lower noise levels both visually and quantitatively, and in-depth analysis shows it acquires data more faithfully. We also demonstrate the combination of the cost-effective CMOS camera with a bare-bones, free-standing microscope system, which allows for super-resolution live-cell imaging at a total cost of less than 20.000 Euro. We aim to contribute to the growing efforts of ‘democratizing’ super-resolution microscopy and allows for a broader applicability of SOFI imaging in the scientific community.

## 4 Materials and Methods

The microscope systems and corresponding hardware that we used for our measurements are described below. For all data acquisition we used the open-source and freely available Micro-Manager software^24,25^ and a custom device adapter providing full speed for the IDS CMOS camera^5,26^. For SOFI data reconstruction we made use of the Localizer^22^ package.

### 4.1 Scientific microscope with 50/50 beam splitter

A *Nikon Eclipse TI2* motorized microscope was used for the direct camera comparison. Excitation light was provided by an *Oxxius L6Cc Laser combiner box* coupled to a *Nikon Ti2-LA-BF* laser illumination unit, providing up to 200 mW excitation light at 488 nm. The sample is imaged through a *CFI Apo TIRF 100XC Oil objective*, excitation and emission light is separated via an AHF / Chroma Technology Corp. ZT405/488/561/640rpcv2 imaging-quality dichroic and matching ZET405/488/561/640 nm emission filters were used.

The emission light forms a 100x magnified intermediate image through the tube lens, present in the microscope. A *f*_1_ = 125 *mm* lens (AC-254-125A, Thorlabs; L2 in figure 1 panel C) is used to re-collimate the light, which then passes the 50/50 beam-splitting cube (ANR: 236513, Qioptiq). It is re-imaged onto the cameras through a second lens, which focal length is varied depending on the camera’s physical pixel size (*f*_2_ = 80 *mm*, 75 *mm*, 200 *mm* for the sCMOS, CMOS and emCCD, respectively; L3 and L4 in figure 1 panel C). In this way, the projected pixel sizes of all three cameras are closely matched to 100 nm (see table 1). The lenses employed here are Thorlabs AC-254-080A, AC-254-075A and AC-254-200A respectively.

The split ratio of the beam-splitting device was tested experimentally, to ensure that any imbalance was small enough to not impact the experiments. For this test, three datasets were analyzed for which the sCMOS camera was mounted on the transmission arm of the beam splitter, and the CMOS camera on the reflective arm, and three datasets where these positions were switched. For each dataset, the average frame brightness (in raw counts) was calculated for both cameras, and added to a scatter plot. A linear fit was performed, and the slopes were compared. Any imbalance between the transmission and the reflection arm would show up in a difference in slope between these configurations. From our data (see fig. 1 panel D) we can verify the imbalance of the splitter to be not more than 1%.

To enable frame-by-frame comparison, an electronic triggering and light source gating system was used. The exposure time and frame rate of the industry-grade CMOS camera can be set independently (introducing non-active delays if the frame rate is lower than the exposure time would allow). This camera was used as a “master clock”, as its exposure output was used as a trigger of either the sCMOS or emCCD camera. As the sCMOS device uses a rolling shutter readout, its *global reset* feature was turned ‘on’ to enable this mode. The excitation laser was gated through its modulation input and was only activated when both cameras were light sensitive. A microcontroller (ATMEL328, Arduino Uno) was used for this signal processing, and the timing scheme was continuously monitored using a digital storage oscilloscope (Rigol DS1054Z).

### 4.2 Home-built microscope with industry-grade CMOS

The home-built system was equipped with a 473 nm, 50 mW diode laser for excitation (Spectra Physics Excelsior, SN 50001, USA). The second laser attached to this system was not used for the experiments. Nevertheless, due to this second laser additional lenses were implemented in the excitation pathway of the 473 nm laser, which would not be necessary if only ffDronpa is used. An acousto-optic tunable filter (‘AOTF’, AA OptoElectronic AOTF, SN 26074 and 25278, France) was used to enable fast switching rates of the laser, only illuminating the sample during data acquisition, reducing photobleaching effects to a minimum. After passing through the AOTF, the excitation beam is coupled into a single mode fiber and then focused into the objective (UPLSAPO60XO 60x, 1.35 NA oil objective, Olympus). The fluorescence signal was collected with the same objective and the emission was separated from the excitation source by a dichroic mirror (HC Dual Line – BS R488/561, Semrock). The emission is then focused on the industry-grade CMOS camera using a 250 mm tube lens and after filtering with an emission filter (593/40 BrightLine HC, Semrock), resulting in an overall projected pixel size of 70 nm.

### 4.3 Transfection and sample preparation

Cos-7 cells were cultured in DMEM supplemented with FBS, glutamax and gentamycin at 37 °C with 5% CO2. Cells were washed and detached from the growth flask using a 0.05% Trypsin solution. The cell suspension was then seeded on 35 mm glass bottom culture dishes (#1.5 thickness, MatTek) to ensure a confluency of 50% to 80% for transfection. Cells were then transfected with pcDNA::MAP4-ffDronpa^18^ using FuGENE6 (Promega) according to manufacturer’s instructions, and cells were incubated for a maximum of 24 hours before imaging. For the imaging process the media was replaced with PBS.

### 4.4 Fourier Ring Correlation

Fourier ring correlation^27–29^ is an established image analysis technique to provide an experimental spatial resolution quantification, without relying on measuring the separation or size of single, isolated features. As input, it requires two images of the same structure, but acquired through independent measurements. This is easy to implement for wide-field, confocal and to some extent SMLM-like techniques, but requires a slightly more involved procedure for SOFI:

To acquire pseudo-widefield input datasets, images 1-5 and 6-10 of each stack were summed up. This proved enough to eliminate artifacts introduced by the blinking of the probes, but less enough to not pick up camera correlation artifacts. These arise when averaging over large amounts of acquired data (e.g. 200 frames of a SOFI stack) and manifest in high-frequency correlations far beyond the microscope’s passband (and thus unphysical). They likely arise due to imperfections in the camera sensor and read-out electronics, giving rise to static noise patterns that yield high-frequency correlations.

To acquire SOFI input datasets, the full stack of 500 images was split in a blocked fashion, i.e. images 1-50, 101-150, …, 401-450 where assigned to dataset A, and 51-100, 151-200, …, 451-500 where assigned to dataset B, and then reconstructed with the SOFI algorithm. This blocking scheme proved necessary, as both of the simpler alternative introduce artifacts: Splitting the stack in the middle (1-250 to A, 251 -500 to B) yields a situation where dataset B is disproportionally affected by photo-bleaching. Splitting the stack in an even/odd fashion (1,3,5, …, 499 to A, 2,4,6, …, 500 to B) avoids this and would be very suitable for localization microscopy, but destroys the time correlation inherent to the data and picked up by the SOFI algorithm.

For consistency with the displayed FRC plots, the SOFI reconstructions shown are the ones that served as input into the FRC analysis, i.e. these images have been reconstructed from 250 frames extracted in 5x 50 blocks of images.

### 4.5 Spatial correlation analysis

The theory for cross-correlation SOFI was introduced in Dertinger et. Al.^30^. The formula given there for second order SOFI can be broken up in three parts: a constant determined by the behavior of the dye, a constant determined by the separation between the pixels used in the calculation and the shape of the PSF and a term depending on the location of the emitters in the sample and the PSF (This final term proves the unbiassed super resolution imaging performance of the technique). To investigate if additional correlations are introduced by the instrument, we want to compare the second constant (determined by the emitter separation) with the theoretical proposed form. Here, we assume that other sources of correlation would not show the same dependency. To make this comparison we use the fact that on average the structure being imaged should be uncorrelated to the camera at a sub-pixel level. Using this information, when for a given shift we average over enough unrelated virtual SOFI pixels, the sample dependent term becomes constant compared to other shifts. In practice, the full SOFI image is calculated for all shifts within a 3 by 3 grid. Afterwards, for each shift, we average the image to yield a single value. According to theory, to a very good approximation the dependence of this value on the distance between the pixels is described by a gaussian curve^30^ where the width of the gaussian is 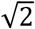 larger than the PSF width. In practice we see that some points follow this curve well, while some do not. The assumption at this point is that the points deviating from this behavior are being caused by additional correlation between the pixels, deriving from a source beside single molecule fluctuations. In the analysis we label these points as ‘spurious correlation’ which we attribute to the camera operation.

## 5 Acknowledgements and author contributions

R.V.D.E. build the beam-splitting system, performed measurements and SOFI reconstructions, and wrote parts of the manuscript. A.S. performed measurements, built the freestanding microscope system, and wrote parts of the manuscript. W.V. performed SOFI reconstructions and analysis. S.D. provided the ffDronpa constructs and helped with the measurements. W.H. performed the cell transfections. P.D. and T.H. conceived of and supervised the project, and helped writing the manuscript. M.M. supervised the projects, performed data analysis, and helped writing the manuscript. All authors participated in the reading and editing of the final manuscript. We would like to thank Mario Lachetta, Marcel Peplonski and Andreas Markwirth for help with and discussion on the opto-mechanical design and the control electronics of the free-standing microscope system. This project has received funding from the European Union’s Horizon 2020 research and innovation program under the Marie Skłodowska-Curie grant agreements No. 752080 and No. 766181. P.D. acknowledges support by the European Research Council via ERC Starting Grant 714688 and from the Research Foundation-Flanders (FWO) via grants G062616N, G0B8817N, G0A5817N, and VS.003.16N. R.V.D.E. thanks the FWO for a doctoral fellowship, S.D. thanks the FWO for a post-doctoral fellowship. There are no competing interests. Raw image data is made available upon request.

